# Systematic analysis of in-source modifications of primary metabolites during flow-injection time-of-flight mass spectrometry

**DOI:** 10.1101/2022.09.28.509873

**Authors:** Niklas Farke, Thorben Schramm, Andreas Verhülsdonk, Hannes Link

**Affiliations:** Bacterial Metabolomics, CMFI, University Tübingen, Auf der Morgenstelle 24, 7206 Tübingen, Germany

**Keywords:** Flow-injection mass spectrometry, Metabolomics, Electrospray ionization, Feature network, In-source modifications

## Abstract

Flow-injection mass spectrometry (FI-MS) enables metabolomics studies with a very high sample-throughput. However, FI-MS is prone to in-source modifications of analytes because samples are directly injected into the electrospray ionization source of a mass spectrometer without prior chromatographic separation. Here, we spiked authentic standards of 160 primary metabolites individually into an *Escherichia coli* metabolite extract and measured the thus derived 160 spike-in samples by FI-MS. Our results demonstrate that FI-MS can capture a wide range of chemically divers analytes within 30 seconds measurement time. However, the data also revealed extensive in-source modifications. Across all 160 spike-in samples, we identified significant increases of 11,013 ion peaks in positive and negative mode combined. To explain these unknown *m/z* features, we connected them to the *m/z* feature of the (de-)protonated metabolite using information about mass differences and MS2 spectra. This resulted in networks that explained on average 49 % of all significant features. The networks showed that a single metabolite undergoes compound specific and often sequential in-source modifications like adductions, chemical reactions, and fragmentations. Our results show that FI-MS generates complex MS1 spectra, which leads to an overestimation of significant features, but neutral losses and MS2 spectra explain many of these features.

**Highlights:**

- FI-MS enables measurements of chemically divers metabolites.
- Extensive in-source modifications during electrospray ionization are detected by FI-MS.
- A network approach explains 49 % of all recorded in-source modifications.

## 1. Introduction

Flow injection mass spectrometry (FI-MS) does not rely on chromatographic separation of analytes [1,2]. Instead, samples are injected into the mobile phase that directly enters a mass spectrometer. Metabolites are then distinguished solely by their mass to charge ratio (*m/z*) in the MS1 spectrum. This makes FI-MS faster than methods with chromatographic separation [3,4], and enables run times on the second-time scale or even real-time metabolomics with living cells [5].

FI-MS has been applied to measure the metabolome in various organisms including *Escherichia coli*, yeast, ruminants, and human cancer cell lines [6–10]. In these studies, hundreds or even thousands of strains or conditions could be analyzed due to the fast measurement time of FI-MS. Although FI-MS detects usually a very large number of *m/z* features (ion peaks in the MS1 spectrum), only a small fraction of *m/z* features can be annotated to metabolites. Thus, there is a large number of unexplained *m/z* features in FI-MS analyses, which could mean that either many metabolites are not known or that single metabolites produce multiple *m/z* features.

Annotation of unknown *m/z* features is a general challenge in all untargeted metabolomics methods [11–14]. For example, an untargeted LC-MS analysis suggested that out of 25,000 measured *m/z* features less than 1,000 originated from unique metabolites [15]. The high number of *m/z* features in untargeted metabolomic methods is often attributed to contaminants, isotopes, modification of metabolites in the ion-source, and other mass spectrometry artifacts.

In-source fragmentation is one example of such mass spectrometry artifacts that increase the number of *m/z* features per metabolite. The conditions in the ESI can lead to fragmentation because metabolites are subjected to high temperatures (150°C to 400°C) and electric potentials between 2000 V and 4000 V. While ESI sources are usually designed to minimize in-source fragmentation, it is also possible to promote in-source fragmentation such that MS1 spectra resemble MS2 spectra that were obtained by collision induced dissociation [16]. Apart from in-source fragmentation, other modifications of metabolites in the ion-source includes the formation of adducts (e.g. with Na, K, ammonia, sulfate), gains or losses of functional groups by chemical reactions (methylation, phosphorylation), or formation of homo- and heterodimers. Even self-cyclization has been observed for glutamate and glutamine [17].

A common approach to identify in-source modifications and improve *m/z* feature annotation is based on chromatographic peak shape correlation analysis [18–20]. This approach considers that *m/z* features from the same metabolite must have the same elution profile [21]. Chromatographic peak shape correlation analysis is especially effective if it is combined with MS2 spectra [19,22,23] or isotope labeled substrates [11,21]. Some recent molecular networking approaches [19,23,24] combine similarities of elution profiles and MS2 spectra to identify in-source modifications and to increase annotation confidence. In isotope labelling approaches, metabolites are labelled by feeding cells with ^13^C-carbon or ^15^N-nitrogen sources [11,21], which changes the mass of all metabolites (N- or C-containing) but not their retention times. Analyzing the mass differences of *m/z* features with the same retention time can then improve annotation confidence and identification of in-source modifications or contaminants.

Because FI-MS lacks a chromatographic separation, it is not possible to detect in-source effects by chromatographic peak shape correlation analysis. Therefore, approaches to consider in-source effects in FI-MS are limited and currently based on extending the list of reference masses [1,25]. This means that, instead of annotating *m/z* features only to (de-)protonated metabolites, they are also annotated to the most prevalent adducts and neutral losses or gains. However, this approach cannot identify complex sequential in-source modifications due to combinatorial explosion of the reference list. Moreover, it is difficult to unequivocally annotate *m/z* features to a single entry in a reference list, especially if they include a large number of metabolites and derivatives with the same mass.

Here, we used an experimental approach to identify in-source modifications of metabolites in FI-MS. Therefore, we spiked 160 metabolite standards individually into an *E. coli* extract and measured MS1 spectra by FI-MS. We then searched for *m/z* features that increased in a spike-in sample relative to all other spike-in samples. On average 68 *m/z* features increased per spike-in standard suggesting extensive in-source modifications. While some spike-in standards showed hundreds of significant *m/z* features that should all originate from a single metabolite standard, others showed only increases of the *m/z* feature that matched the (de-)protonated metabolite standard. We could explain 49% of the significant *m/z* features by connecting them in networks that represent known in-source reactions, adducts, isotope patterns, and in-source fragments.

## 2. Materials and Methods

### 2.1 Chemicals and materials

Authentic metabolite standards were purchased from Merck KGaA (former Sigma-Aldrich, Germany). The standards were dissolved in water to a concentration of 1 mM if not stated otherwise (**Supplementary Data: Table A**). Standards were then further diluted with acetonitrile and methanol to a final concentration of 10 μM in 40:40:20 acetonitrile:methanol:water. The 10 μM metabolite standards were then added to an *E. coli* metabolite extract to yield a final concentration of 1 μM. *E. coli* cultures were in a M9 minimal medium, which contained: 22 mM KH_2_PO_4_, 42.2 mM Na_2_HPO_4_, 11.3 mM (NH_4_)_2_SO_4_, 8.56 mM NaCl, 100 μM CaCl_2_ x 2 H_2_O, 1 mM MgSO_4_ x 7 H_2_O, 60 μM FeCl_3_, 6.3 μM ZnSO_4_ x 7 H_2_O, 7.6 μM CoCl_2_ x 6 H2O, 7.1 μM, 7 μM CuCl_2_ x 2 H_2_O, 2.8 μM Thiamine-HCl, and MnSO_4_ x 2 H_2_O.

### 2.2 Metabolite extracts from *E. coli* cultures

5 mL LB medium was inoculated with *E. coli* MG1655 from a cryo stock. After 6 - 7 h of cultivation at 37°C, 10 μL of the culture was transferred to 5 mL M9 minimal medium with 5 g/L glucose. For ^13^C-labelled extracts, uniformly labelled ^13^C-glucose was used (#CLM-1396, Cambridge Isotope Laboratories Inc., USA). The M9 precultures were grown overnight at 37°C and at 220 rpm shaking. 20 mL of M9 with 5 g/L ^12^C- or ^13^C-glucose was inoculated with the overnight culture to an optical density at 600 nm (OD) of 0.05. At an OD of 1, aliquots of 4 mL of the culture were vacuum-filtrated using 0.45 μm pore size filters (HVLP02500, Merck Millipore). The filters were transferred to -20°C cold 40:40:20 acetonitrile:methanol:water for metabolite extraction. After at least 30 min at -20°C, the metabolite extracts were centrifuged for 30 min at -9°C and 4,000 rpm. The supernatant was stored at -80°C.

### 2.3 Mass spectrometry

Samples were analyzed by FI-MS on an Agilent 6546 Series quadrupole time-of-flight mass spectrometer (Agilent Technologies, USA). The electrospray source was operated in negative and positive ionization mode. The mobile phase was 60:40 isopropanol:water buffered with 10 mM ammonium carbonate (NH_4_)_2_CO_3_ and 0.04 % (v/v) ammonium hydroxide for both ionization modes, and the flow rate was 0.15 mL/min. For online mass axis correction, 2-propanol (in the mobile phase) and HP-921 were used for negative mode and purine and HP-921 were used for positive mode. Mass spectra were recorded in profile mode from 50 to 1100 *m/z* with a frequency of 1.4 spectra/s for 0.5 min using 10 Ghz resolving power. Source temperature was set to 225 °C, with 1 L/min drying gas and a nebulizer pressure of 20 psi. Fragmentor, skimmer, and octupole voltages were set to 120 V, 65 V, and 650 V, respectively.

### 2.4 Data preprocessing

Raw files were converted into “mzXML” format by MSConvert [26]. Further data processing was performed using MATLAB version R2021a (The Mathworks, Inc., USA). For each sample, an average spectrum was calculated from the ten scans with the highest ion counts. The spectra were resampled to 10^6^ data points to align *m/z* values of all samples. Ion peaks were picked with the “findpeaks” function of MATLAB, using a peak height and prominence cutoff of 1,000 units. Hierarchical clustering with a tolerance of 7.5 mDa was used to bin peaks. For each peak bin, we calculated a centroid *m/z* value from the individual peak *m/z* values. Peaks were annotated to metabolites using the centroid *m/z* value with a tolerance of 3 mDa. Z-scores were calculated from logarithmic mean signal intensities (triplicates). Z-scores above three were considered significant.

### 2.5 Calculation of mass differences

Mass differences between all significant features were calculated and combined for positive and negative ionization mode. Using the MATLAB function “histcounts”, all mass differences were assigned to one of 5 × 10^5^ bins, which accounted for a mass resolution of ca. 2 mDa. The total number of mass differences in each bin is the frequency of a mass difference. In the resulting neutral loss spectrum (x-axis is the mass difference and y-axis the frequency), peaks were picked with 3 mDa tolerance using the “findpeaks” function with prominence and height cutoffs of 10 units. 51 peaks that matched mass differences in the literature [23] were then used for further analysis (**Supplementary Data: Table B**).

### 2.6 Construction of feature networks

Networks of significant *m/z* features were constructed for each spike-in sample in positive and negative ionization mode. Nodes are significant *m/z* features, and edges are putative modifications like adducts or chemical reactions, or isotopes. Edges are drawn between each pair of nodes if the mass difference between them matches the mass differences in the list of 51 explained mass differences (**Supplementary Data: Table B**). The fraction of explained features are calculated by counting all features that are either directly or indirectly connected to the (de-)protonated metabolite or fragment ion versus the total number of significant features for each spike-in standard. The feature networks were built with python v. 3.8.5 using the “networkx” toolbox. MS2 spectra were obtained from the Human Metabolome Database when experimental spectra were reported for a spike-in standard [27] (**Supplementary Data: Table C**). The experimental MS2 spectra from HMDB contained data that was acquired by high- and low-resolution mass spectrometers. Therefore, we matched our significant *m/z* features to the MS2 spectra with a tolerance of 100 mDa.

## 3. Results

### 3.1 FI-MS with 160 authentic metabolite standards

We prepared 160 authentic standards of primary metabolites and spiked them individually into a metabolite extract from glucose-fed *E. coli* cells (**Supplementary Data: Table A**). The 160 standards fall into six functional categories: amino acid metabolism, nucleotide metabolism, central metabolism, cofactor metabolism, antioxidants, and others. Each metabolite standard was added to the *E. coli* metabolite extract at a final concentration of 1 μM and was measured by FI-MS in both positive and negative ionization mode (three analytical replicates) (**Figure 1a**).

**Figure 1.**
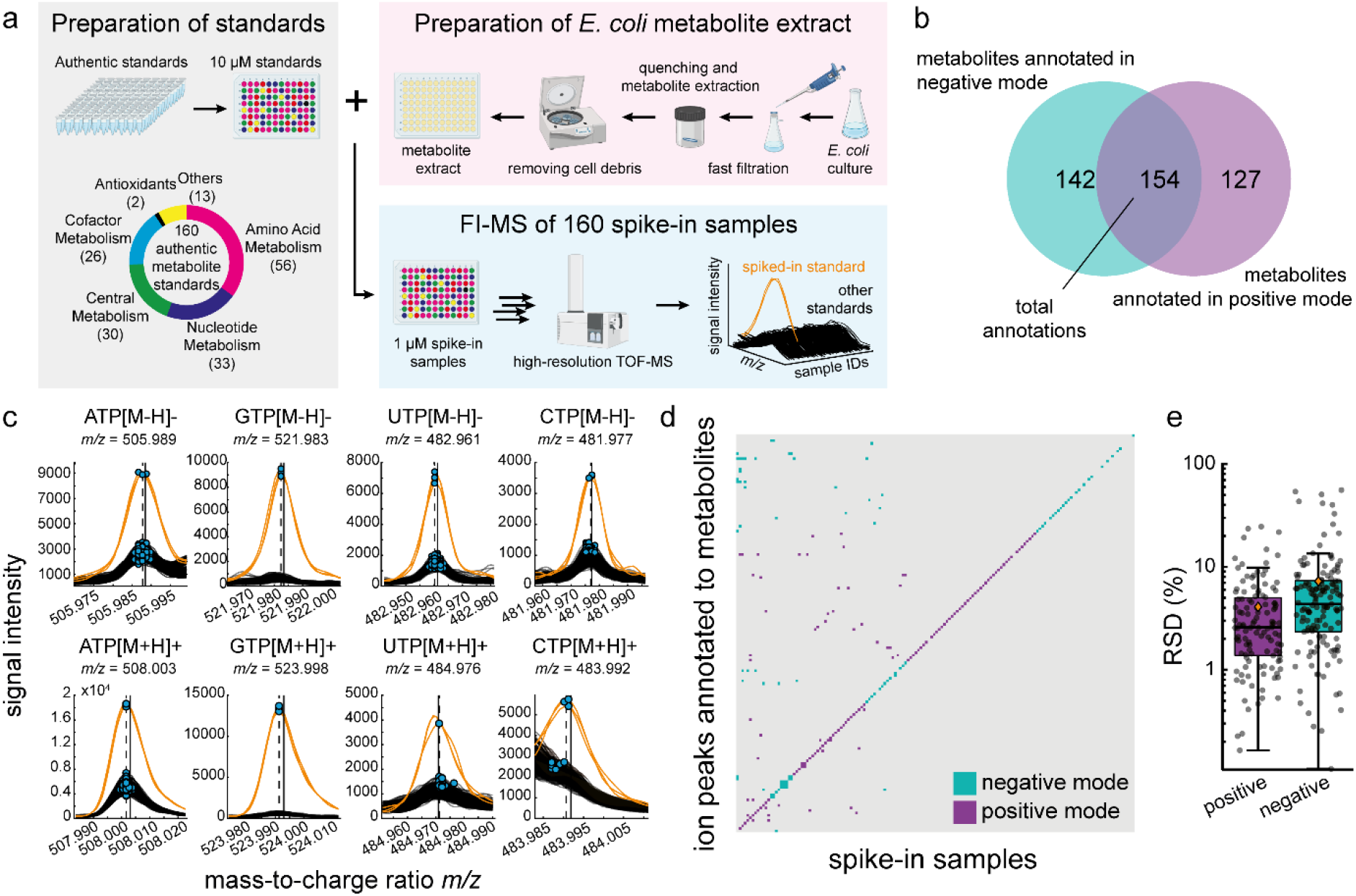
FI-MS with 160 authentic metabolite standards. a) 160 authentic metabolite standards were spiked into an *E. coli* metabolite extract at a final concentration of 1 μM. Spike-in samples were measured by FI-MS in analytical triplicates in positive and negative ionization mode. b) Number of ion peaks (*m/z* features) that were annotated to the protonated (positive mode) and deprotonated (negative mode) form of the 160 metabolite standards. c) Ion peaks that are annotated to four nucleotides (ATP, CTP, GTP, UTP) in positive and negative ionization mode. The spike-in sample that contains the respective nucleotide is indicated in orange, the other 159 spike-in samples are black. Blue dots indicate *m/z* features in single samples, and vertical dotted lines are merged and centroided *m/z* features. Vertical solid lines indicate the monoisotopic masses of the nucleotides plus/minus the mass of a proton. d) The binary heatmap shows increases of *m/z* features that are annotated to metabolite standards in the spike-in samples. Significant increases of *m/z* features (z-score > 3) are shown in blue (negative mode) and purple (positive mode). Columns are the spike-in samples and rows the respective *m/z* features. e) Boxplots show the relative standard deviation (RSD) of metabolite standards. Black dots are the RSD for each spike-in metabolite (n = 3). Orange diamonds are the means.

Out of 160 metabolite standards, 154 were annotated in their protonated or deprotonated form to an ion peak in the MS1 spectrum (**Figure 1b**). Six metabolites were not annotated, either due to low abundant ion peaks (< 1,000 counts: menadione, 3,4-dihydroxy-L-phenylalanine, tetrahydrofolic acid, carbamoyl-P, and L-cysteine) or because the ion peak prominence was too low (< 1,000 counts: argininosuccinic acid).

Next, we inspected if the addition of a metabolite standard led to increases of the respective ion peak. For example, spike-in samples with ATP, GTP, CTP, and UTP showed increases of ion peaks that matched the protonated and deprotonated form of these metabolites (**Figure 1c**). Notably, increases of all nucleotides were consistent between three analytical replicates, showing that FI-MS is reproducible. Ion peaks of ATP, GTP, CTP, and UTP were also present in the other spike-in samples (black spectra in **Figure 1c**), but the corresponding ion intensities were often low and close to the baseline signal. These “near-baseline” ion peaks of ATP, GTP, CTP, and UTP were not present in a ^13^C-labeled *E. coli* extract, thus confirming that these peaks originate from endogenous *E. coli* nucleotides (**Supplementary Data: Figure S1**).

In 134 spike-in samples, the ion peaks that matched the (de-)protonated metabolites were significantly increased in either ionization mode (z-score > 3, **Figure 1d, Supplementary Data: Table D**). In negative ionization mode, 120 spike-in samples showed increased signals as deprotonated metabolites ([Metabolite-H]-). In positive ionization mode, 105 peaks increased as protonated metabolites ([Metabolite+H]+). In the following, we will refer to significantly changed ion peaks with a z-score >3 as “significant features”. 26 spike-in standards did not show a significant feature at the (de-)protonated ion peak. One explanation for this is that the metabolites have already a high concentration in the *E. coli* metabolite extract and that an addition of 1 μM does not lead to a strong increase with a z-score > 3. For example, reduced glutathione is one of the most abundant metabolites in *E. coli* [28], and the addition of glutathione standard hardly increased its concentration in the spike-in sample (**Supplementary Data: Figure S2**).

FI-MS was reproducible because the median relative standard deviation (RSD) between the three analytical replicates was below 5 % for signals from metabolite standards in negative and positive ionization mode (**Figure 1e**). Signals from endogenous metabolites, had a median RSD of 12.4 % in positive and 19.4 % negative mode (**Supplementary Data: Figure S3**).

These results suggested that FI-MS can detect concentration changes of chemically diverse metabolites, which are in the physiological range of intracellular metabolites (1 μM in the final sample corresponds to ca. 1 mM intracellularly). However, we noticed that many significant features did not match the (de-)protonated form of the metabolite in the spike-in sample (e.g. significant features that are off the diagonal in **Figure 1d**). Thus, we next investigated all significant features in all spike-in samples.

### 3.2 Single metabolites can produce extensive in-source derivates

Most spike-in samples showed significant features (ion peaks with a z-score > 3) that matched the protonated or deprotonated metabolite (**Figure 1d**). However, most spike-in samples had more significant features than only the (de-)protonated metabolite standard (**Figure 2a**). On average, we found 68 significant features per spike-in sample, and 11 spike-in samples showed more than 100 significant features. The glycerol 3-phosphate(G3P) spike-in sample had the highest number of significant features (5,464, **Figure 2a**). Across all 160 spike-in samples, FI-MS in positive mode showed 10,206 significant features and 807 in negative mode (**Supplementary Data: Table E**). The significant features were distributed over the entire mass spectrum and occurred even in the higher mass range of 800 – 1,000 *m/z* (**Figure 2b**).

**Figure 2.**
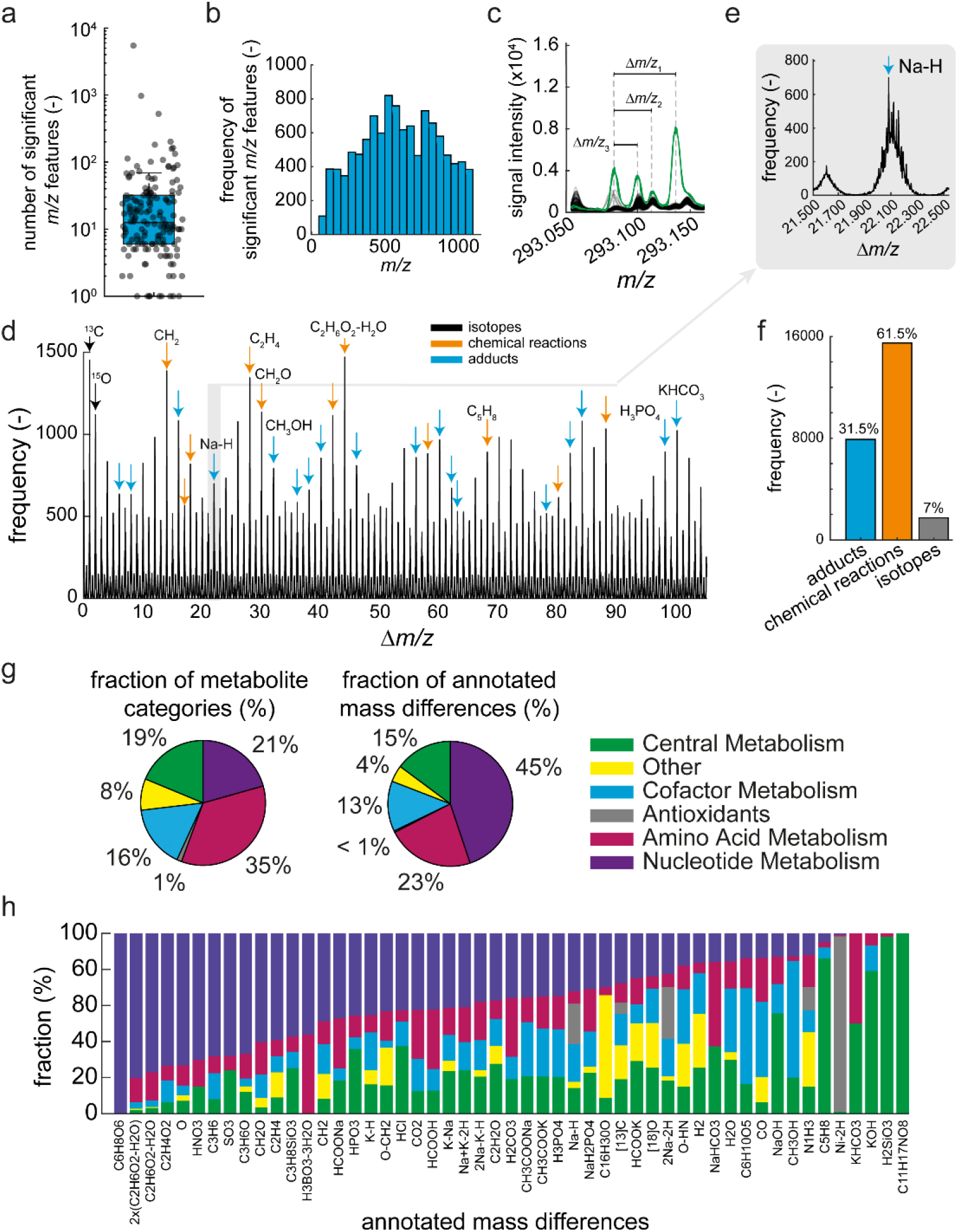
Systematic analysis of all *m/z* features that increase in spike-in samples. a) Number of *m/z* features with a z-score > 3 (significant *m/z* features) in each of the 160 spike-in samples (grey dots). The upper and lower edges of the box in the boxplot indicate the 25 % and 75 % percentiles, and the line is the median. b) Histogram showing the distribution of significant *m/z* features over the MS1 spectrum. c) Example of significant *m/z* features in the MS1 spectrum (100 mDa window) of the spike-in sample with glycerol 3-phosphate. Green lines are the glycerol 3-phosphate spike-in samples (n = 3), and black lines are the other spike-in samples. Rulers indicate the mass differences between two *m/z* features in the spectra. d) Δ*m/z* spectrum based on the pairwise mass differences between all significant *m/z* features in all 160 spike-in samples (shown is the Δ*m/z* range between 0 Da and 110 Da). The peak height corresponds to the frequency of a Δ*m/z* value. Arrows indicate Δ*m/z* peaks that match mass differences of known isotopes, chemical reactions or adducts (**Supplementary Data: Table B**). e) Example of the Δ*m/z* peak that matches the sodium adduct [Na-H]. f) Fraction of Δ*m/z* peaks that match known isotopes, chemical reactions or adducts. g) The left pie chart shows the fraction of metabolite categories across the 160 standards. The right pie chart shows the fraction of annotated mass differences for each metabolite category. h) Stacked bar plot showing the relationship between the functional categories of the spiked-in metabolites and the annotated mass differences. The fraction indicates the ratio between the number of spike-in metabolites of a specific category, in which the mass difference occurred, and the total number of samples, in which the mass difference occurred. The spike-in samples of glycerol 3-phosphate and fumarate were left out.

To understand the origin of these significant features, we first calculated the mass differences (Δ*m/z*) between all pairs of significant features in a single spike-in sample (**Figure 2c**). Several mass differences (Δ*m/z*) occurred frequently across the 160 spike-in samples, thus, indicating common in-source effects like neutral losses, adduct formation, and chemical reactions that are prevalent for many different compounds (**Figure 2d and 2e**). 51 mass differences that appeared more than ten times matched known in-source effects and isotope pattern reported in the literature [23] (**Supplementary Data: Table B**). Out of these 51 known mass differences, 23 were chemical reactions, 26 were adducts, and 2 were natural isotopes (^13^C and ^18^O). The 23 chemical reactions account for 61.5 % of the frequent mass differences (>10 times in all samples), the 22 adducts for 31.5%, and the isotopomers containing ^13^C or ^18^O for 7 % (**Figure 2f**). For example, the 21.982 Da mass difference of a Na-H neutral loss occurred in total 699 times and was among the most frequent ones (**Figure 2e**). The ten most frequent mass differences occurred more than 1,000 times across all 160 spike-in samples, and eight of them could be explained with the mass differences in the literature (**Figure 2d**).

We then wondered whether certain mass differences occurred more frequently for metabolites of a specific functional category than for metabolites of other categories. For example, only 21 % of the 160 standards were metabolites from nucleotide metabolism. Yet, they accounted for 45 % of all explainable mass differences (**Figure 2g**). This indicated that metabolites from nucleotide metabolism were more susceptible to modifications than metabolites in other categories. In contrast, 35 % of the 160 metabolite standards were part of amino acid metabolism but they covered only 23 % of the explainable mass differences indicating that metabolites from amino acid biosynthesis were less prone to modifications in our reference list of mass differences than the other metabolites. Metabolites from central metabolism as well as cofactor biosynthesis accounted for 19 % and 16 % of the standard library, respectively, and they explained a similar fraction of mass differences (15 % and 13 %).

Since some metabolite categories were more often modified than others, we looked into individual mass differences and examined whether certain mass differences occurred preferably for specific categories (**Figure 2h**). Indeed, the data indicated that individual mass differences were more frequent for some categories than for others. For example, the C_6_H_8_O_6_ neutral loss occurred exclusively in metabolites from nucleotide metabolism. Similarly, many other modifications, including modifications with O, HNO_3_, C_2_H_4_O_2_, or SO_3_, occurred more frequently with metabolites from nucleotide metabolism (**Supplementary Data: Table F**).

### 3.3 A network approach explains significant *m/z* features in FI-MS spectra

We expected that the significant features can be linked to the (de-)protonated spike-in metabolite by single and multiple modification steps. To test this, we created a network for each spike-in sample, in which nodes represent all significant features. Then, we drew an edge between two nodes if the mass difference Δ*m/z* between them matched one of the 51 frequent mass differences (the mass differences identified above, see **Figure 2h**). Thus, edges represent in-source effects and nodes significant features (see schematic in **Figure 3a**).

**Figure 3.**
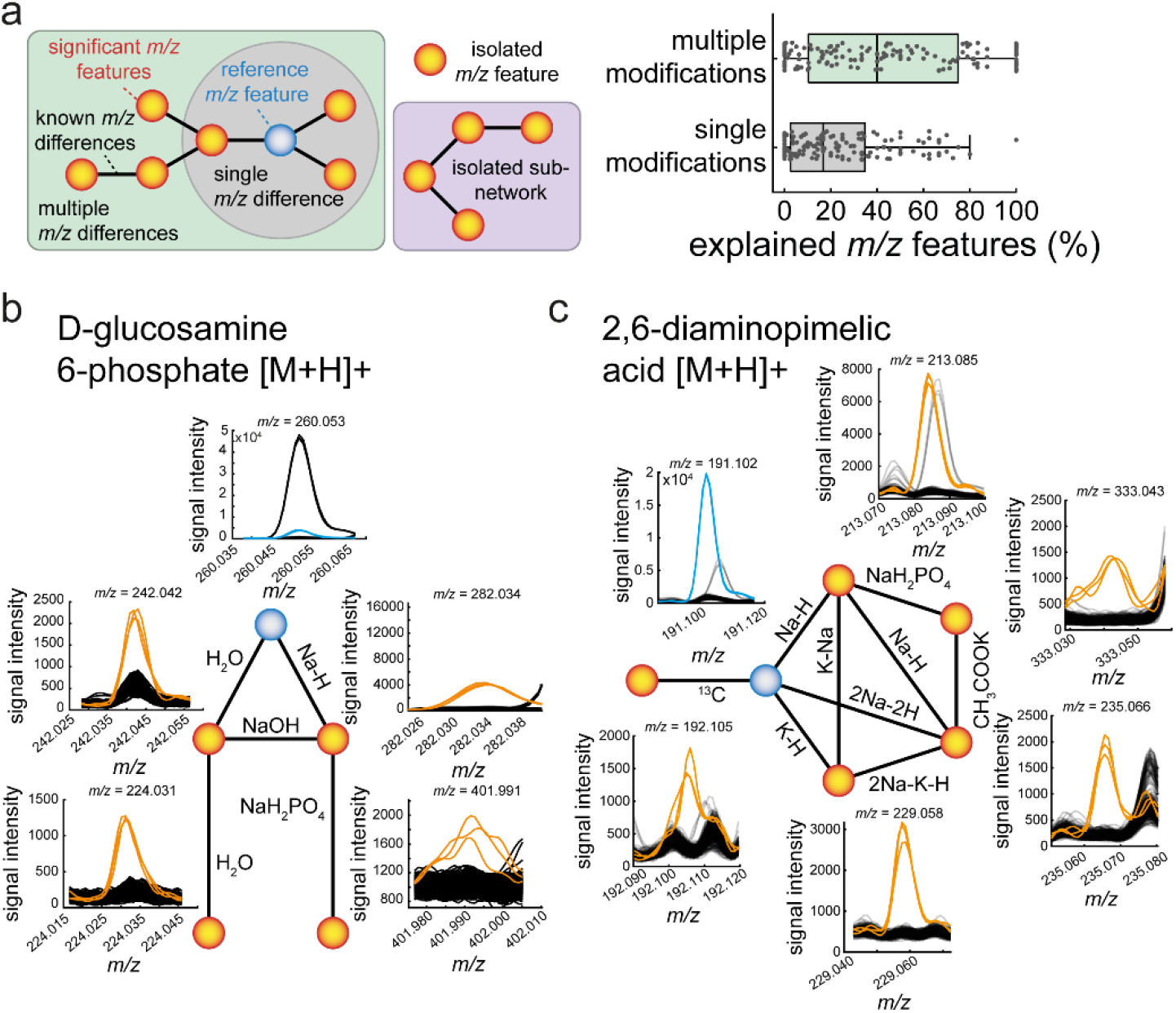
Networks of significant m/z features. a) Concept figure showing the structure of a *m/z* feature network: orange nodes are significant *m/z* features. The blue node is a significant *m/z* feature that is annotated to the (de-)protonated metabolite standard (reference *m/z* feature). Nodes are connected by edges that correspond to one of 51 known *m/z* differences. The grey circle indicates nodes that are directly connected to the reference *m/z* feature by a single *m/z* difference. The green box indicates a network, in which all nodes are connected with the reference node including multiple sequential combinations of *m/z* differences. Isolated subnetworks are not connected to the reference node (purple box). Isolated *m/z* features are not connected to any other node. The boxplots show the fraction of significant *m/z* features that are connected to the reference *m/z* feature by a single *m/z* difference or by multiple sequential combinations of *m/z* differences. b) Example of the feature network of the spike-in sample with D-glucosamine 6-phosphate (Ga6p). c) Same as b) for the diaminopimelic acid (DAP) spike-in sample in positive ionization mode.

The thus derived networks connected on average 43 % of the significant features in a spike-in sample to the *m/z* feature of the respective (de-)protonated metabolite (**Figure 3a**). Thus, 43 % of the significant features can be linked to a single metabolite and therefore, are explained by the 51 frequent mass differences. Only 20 % of the significant features were directly linked to the *m/z* feature of the (de-)protonated metabolite (**Figure 3a, Supplementary Data: Table G**). This shows that single in-source modifications account only for half of the significant features and that sequential modifications are frequent.

For example, the glucosamine 6-phosphate (Ga6p) spike-in standard showed five significant features, which are all directly or indirectly connected to the *m/z* feature that matches protonated Ga6p (**Figure 3b**). Two significant features are directly connected to the protonated Ga6p mass, and they are likely a water loss (H2O) and a sodium adduct (Na-H). Two other significant features (*m/z* = 224.031 and *m/z* = 242.042) were two steps away from protonated Ga6P, and they were explained by a double loss of water and a NaH_2_PO_4_ adduction to the sodium adduct. Thus, drawing edges in an unbiased way between all pairs of nodes resulted in a network that explained all significant features of the Ga6p spike-in sample.

In many networks, the nodes (significant *m/z* features) were connected to the (de-)protonated spike-in metabolite by different series of sequential modifications. One specific series of sequential modifications could be an initial modification by Na-K that is followed by a second modification like Na-H. In some cases, different series of sequential modifications have very similar net mass changes and can explain the same significant feature. One example that illustrates this phenomenon is the feature network of 2,6-diaminopimelic acid (DAP) in positive ionization mode. The DAP spike-in sample showed six significant features, which were all connected with the *m/z* feature of protonated DAP (**Figure 3c**).

The network approach connected on average 43 % of all significant features of a spike-in sample to the (de-)protonated metabolite standard. Yet, some significant features had no connection to others or they formed sub-networks with no connection to the (de-)protonated metabolite (see schematic in **Figure 3a**). Therefore, we next examined whether sub-networks and isolated features are caused by in-source fragmentation, which can lead to similar effects as collision induced dissociation in tandem mass spectrometry [16].

### 3.4 MS2 information identifies significant features that are in-source fragments

To identify significant features that originate from in-source fragmentation of the metabolite standard, we used information about MS2 spectra in the human metabolome database (HMDB [27]). HMDB listed experimental MS2 spectra for 152 out of 160 metabolite standards (**Figure 4a, Supplementary Data: Table C**). 103 standards had at least one significant feature that matched an ion peak in the MS2 spectrum (**Supplementary Data: Table H**). On average, each spike-in sample had 3.4 in-source fragments indicating a substantial number of in-source fragmentation events during FI-MS (**Figure 4a**). In total, 551 MS2 features matched the significant *m/z* features. As expected, in-source fragments had masses in the lower *m/z* range, between 50 and 500 *m/z* (**Figure 4b**). One example of an in-source fragment is hypoxanthine, which is formed by fragmenting inosine and inosine monophosphate (IMP). Consequently, the ion peak of hypoxanthine increased in the IMP and inosine spike-in samples (**Figure 4c**).

**Figure 4.**
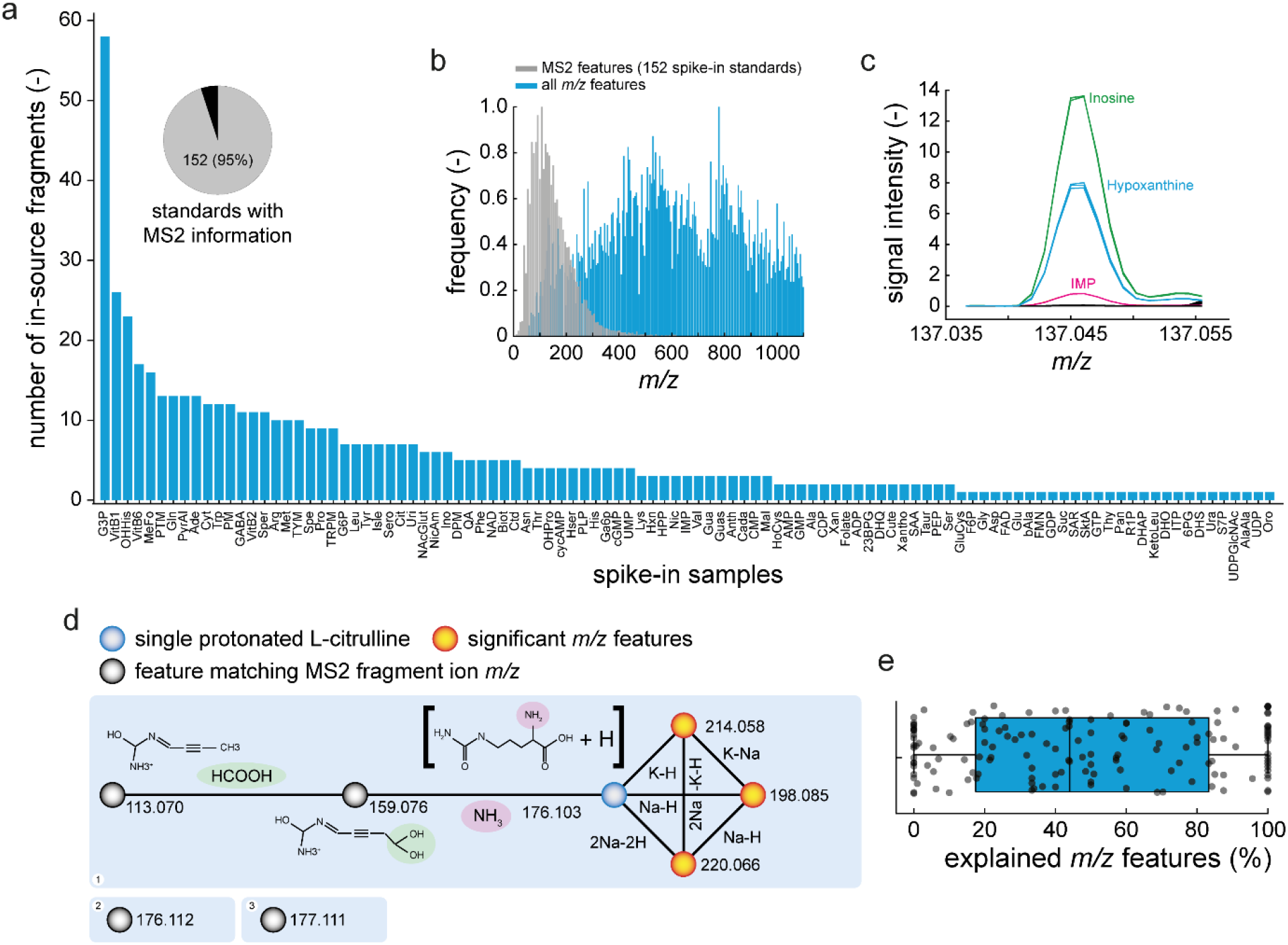
Identification of in-source fragments by MS2 spectra of metabolite standards. a) The pie chart shows the fraction of metabolite standards with MS2 information in the human metabolome database (HMDB). The bar plot shows the number of MS2 fragment masses that match the features of individual spike-in samples. b) Histogram showing the distribution of the *m/z* features. All significant *m/z* features from the spike-in samples are blue. MS2 fragment masses are grey. c) Example MS1 spectrum at the mass of hypoxanthine. The purple line is the IMP spike-in sample, the blue line is the hypoxanthine spike-in sample, and the green line is the inosine spike-in sample. d) Example network of L-citrulline. Nodes are features and edges are explained mass differences. Black nodes are features that matched MS2 fragment masses from HMDB. The blue node is the protonated mass of citrulline. The orange nodes are other features. For the features at *m/z* = 113.070 and *m/z* = 159.076, structures were predicted by CFM-ID. e) Boxplot showing the fraction of explained features for all spike-in samples. Each black point corresponds to a spike-in sample and shows the explained fraction of features. Upper and lower box edges indicate the 25 % and 75 % percentiles. The whiskers indicate the furthest point, at which samples were not considered as outliers. The black line indicates the median.

Then, we tested whether fragments were present in sub-networks without a link to the metabolite standard. For example, the L-citrulline spike-in sample showed 8 significant features, 6 of which were connected with the *m/z* feature of the (de-)protonated metabolite (**Figure 4d**). Two significant features were isolated nodes with no connection, but they were in the MS2 spectrum of citrulline (*m/z* = 177.111 Da and *m/z* = 176.112 Da). The MS2 spectrum included another two *m/z* features that were already linked to the metabolite standard (*m/z* = 113.070 Da and *m/z* = 159.076 Da), thus indicating that collision induced dissociation (CID) produces some of the 51 in-source modifications. In case of L-citrulline, these were a neutral loss of NH3 and a neutral loss of a HCOOH group. To confirm that these losses occur by CID, we predicted fragment structures of L-citrulline by CFM-ID [29]. Indeed, CFM-ID predicted the fragment structures that matched the masses of the two *m/z* features (113.070 Da and 159.076 Da) and confirmed the neutral losses of NH3 and HCOOH (**Figure 4d**).

In total, adding in source fragments to our networks explained another 6 % of significant *m/z* features. Thus, on average, 49 % of all significantly changed *m/z* features were connected with the correct metabolite standard either by known in-source modifications or by in-source fragmentation (**Figure 4e, Supplementary Data: Table G**).

### 3.5 Misannotation of in-source derivates to metabolites

A single metabolite can produce multiple significant features, and we wondered how many of these significant features were misannotated to a metabolite that was not spiked into the sample. To determine how many significant features were misannotated, we used a reference list of 961 *E. coli* metabolites from the genome-scale metabolic model iML1515 [30]. Since FI-MS cannot resolve isomers, they were considered as a single metabolite.

In 54 % of our standards (87/160), at least one significant feature was falsely annotated to a metabolite (**Figure 5**). 18 % of the standards with misannotations had more than two misannotations. Overall, 64 % of the misannotations were in positive ionization mode (131 in total) and 37 % were in negative ionization mode (75 in total) (**Figure 5, Supplementary Data: Table I**). This means that biological screens with FI-MS are prone to misannotations if only (de-)protonated masses are considered. Based on our results, an estimate is that one (true) increase of a single metabolite will cause one (false) increase of one ion peak that is misannotated to another metabolite.

**Figure 5.**
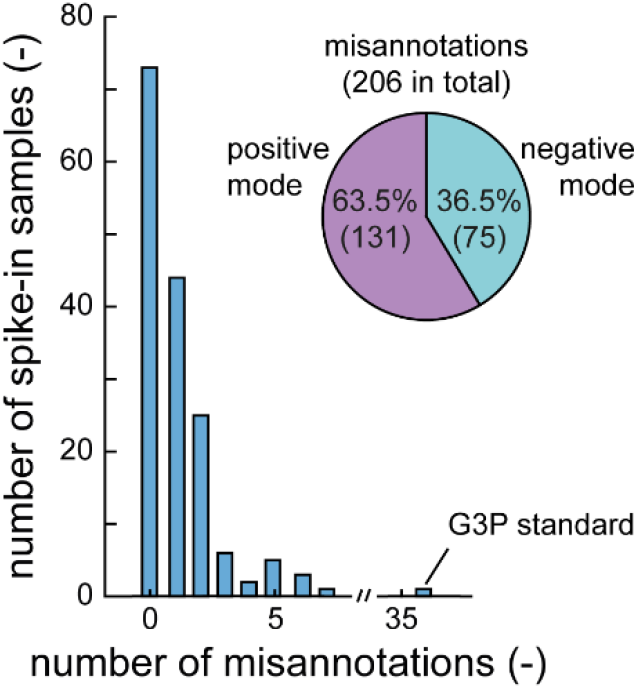
Misannotation of significant features to metabolites. The histogram shows the number of significant features that are misannotated to a metabolite. The pie chart shows the fraction of misannotations for positive and negative ionization mode.

## 4. Discussion

FI-MS methods have been used for metabolome analyses in various studies [1,2,4–10]. Their advantages are fast analysis times (10 to 30 seconds per sample) and a high coverage of metabolites (often more than 1,000 putatively annotated metabolites [6,7]). Disadvantages, however, are low confidence levels of metabolite annotation and a high susceptibility to matrix effects due to the lack of chromatographic separation. Here, we confirmed the broad metabolite coverage of FI-MS, which detected increases of 134 out of 160 metabolite standards based on (de-)protonated ion peaks in the MS1 spectrum. However, we also observed pervasive in-source modification of metabolites. These in-source modifications lead to multiple ion peaks per metabolite in a MS1 spectrum and, in the worst case, to false positive hits in FI-MS analyses of biological samples. By systematically analyzing FI-MS data from 160 spike-in standards, we found that a single metabolite produces on average 68 significant features and that, in extreme cases, more than 1,000 significant features originate from only one metabolite. This observation matches previous LC-MS based studies where the majority of *m/z* features were attributed to in-source modifications and only few (3 – 5 %) *m/z* features were unique metabolites [11,12,15,31].

Chromatographic peak shape correlation of *m/z* features can identify such confounding effects in LC-based methods [19,21], but they are difficult to detect with FI-MS methods. Here, we used an experimental approach and examined significant *m/z* features in metabolite standards, which are most likely in-source derivates. Connecting these features via 51 mass differences of neutral losses, adducts, and isotopes described in the literature [23] resulted in networks that explained the origin of 43 % of the significant features. MS2 spectra of the metabolite standards provided additional information about in-source fragmentation and explained another 6 % of the significant features.

Taken together, we found that FI-MS of single metabolites produces complex MS1 spectra, but they are explainable by known in-source modifications. The jury is out if in-source modifications are a bug or a feature for FI-MS data analysis: they may complicate or improve metabolite annotation. Therefore, the future challenge is to use FI-MS spectra of single metabolites to deconvolute FI-MS spectra from biological samples and, thereby, increase confidence of metabolite annotation. Here we provided a starting point with in-source modifications of 160 metabolites (**Supplementary data: Table E**) and mining MS1 spectra in existing databases like GNPS [32] and METLIN [33] may provide additional reference data.

## Supporting information

Farke_etal_Supplementary_Tables

## Appendix A. Supplementary Data

Supplementary Tables (.xlsx)

Supplementary Figures (.docx):

- Figure S1: Ion peaks of nucleotides in negative ionization mode from fully labelled ^12^C- and ^13^C-labelled *E. coli* extracts.
- Figure S2: Ion peak of reduced glutathione in negative ionization mode.
- Figure S3: Boxplot showing the relative standard deviation (RSD) of the endogenous metabolites measured by FI-MS in positive and negative ionization mode.

## CRediT authorship contribution statement

‡N. F. and T. S. contributed equally. Conceptualization: N.F., T.S., H.L., Data curation: N.F., Formal analysis: N.F., T.S., Funding acquisition: H.L., Investigation: N.F., T.S., A.V., Methodology: N.F., Project administration: H.L., Software: N.F., T.S., H.L., Supervision: H.L., Visualization: N.F., T.S., Writing – original draft: N.F., T.S., H.L. All authors have given approval to the final version of the manuscript.

## Acknowledgements

This work was supported by the ERC starting grant 715650. Figure 1a and the abstract figure were created in part with BioRender.com.

